# Sensory experience selectively reorganizes the late component of evoked responses

**DOI:** 10.1101/2020.09.30.321547

**Authors:** Edgar Bermudez-Contreras, Andrea Gomez-Palacio Schjetnan, Artur Luczak, Majid H. Mohajerani

## Abstract

In response to sensory stimulation, the cortex exhibits an early transient response followed by a late and slower activation pattern. Recent studies suggest that the early component represents features of the stimulus while the late component is associated with stimulus perception. Although very informative, these studies only focus on the amplitude of the evoked responses to study its relationship with sensory perception. In this work we expand upon the study of how patterns of evoked and spontaneous activity are modified by experience at mesoscale level using voltage and extracellular glutamate transient recordings over widespread regions of mice dorsal neocortex. We find that repeated tactile or auditory stimulation selectively modifies the spatiotemporal patterns of cortical activity, mainly of the late evoked response in anesthetized mice injected with amphetamine and also in awake mice. This modification lasted up to 60 minutes and results not only in an increase in amplitude of the late response after repeated stimulation, but also in an increase in the similarity between the spatiotemporal patterns of the late and the early evoked response patterns in anesthetized mice. This similarity increase occurs only for the evoked responses of the sensory modality that received the repeated stimulation. Thus, this selective long-lasting spatiotemporal modification of the cortical activity patterns might provide evidence that evoked responses are a cortex-wide phenomenon. This work opens new questions about how perception-related cortical activity changes with sensory experience across the cortex.

## Introduction

The ability to learn from and adapt to changes in their environments are crucial skills that allow organisms to survive. The neural correlates of such adaptive processes correspond to changes in patterns of brain activity. The study of how these changes are encoded and transformed by the brain is crucial to understand brain computation (Panzeri et al. 2017). One of the main approaches to study the neural coding problem – to understand what features of brain activity encode information about the stimulus – is to analyze the dynamics of evoked brain responses. Recently, this approach has been used to study sensory perception for discrimination tasks in the somatosensory and visual cortices in rodents and humans (Sachidhanandam et al. 2013; Funayama et al. 2015a; Manita et al. 2015a; Yamashita and Petersen 2016). In these experiments, it was shown that the evoked responses have two components, an early and a late evoked deflections. When compared, hit trials, in which the perception of the sensory stimulus is arguably better, show a larger late evoked deflection than in trials where the animals fail to discriminate the stimulus. Moreover, when the late evoked response is inactivated, the task performance decreases (Sachidhanandam et al. 2013; Funayama et al. 2015a). Therefore, these results suggest a causal role of the amplitude of the late evoked response for the performance in sensory discrimination tasks due to sensory perception being affected when the late evoked response is perturbed. In humans, the biphasic structure of the sensory evoked response has been reported and researched extensively using functional magnetic resonance imaging (fMRI) and electroencephalography (EEG) (Dinteren et al. 2014; Hedges et al. 2016) and magnetoencephalography (MEG) to study cognitive functions (Sutton et al. 1965; Otzenberger et al. 2005; Patel and Azzam 2005; Arrubla et al. 2013; Dinteren et al. 2014; Twomey et al. 2015; Baykara et al. 2016).

In addition to sensory perception, the study of evoked responses has also advanced our understanding of learning and brain plasticity (Cooke and Bear 2010; Takeuchi et al. 2014). In fact, pioneer *in vitro* electrophysiological studies of changes in synaptic efficacy are one of the pillars of the current understanding of brain plasticity (Bear and Malenka 1994; Malenka and Bear 2004). More recent studies have demonstrated that experience can modify the temporal structure of firing patterns in different brain regions in freely behaving rodents (Skaggs and Mcnaughton 1996; Ji and Wilson 2007) and even in anesthetized preparations (Han et al. 2008; Bermudez Contreras, Gomez Palacio Schjetnan, et al. 2013). While changes to evoked responses induced by repeated stimulation has been extensively studied in these works, much less in known about how this activity propagates in the cortex. Here we present a study of the changes induced by repeated sensory stimulation to the spatiotemporal dynamics of the sensory evoked responses in different modalities in anesthetized mice injected with amphetamine and awake mice using wide-field voltage-sensitive dye (VSD) and glutamate imaging respectively. We have previously demonstrated that repeated sensory stimulation induces changes in cortical activity during desynchronized states (Bermudez Contreras, Schjetnan, et al. 2013). Therefore, we injected amphetamine to the anesthetized mice to increase the chances of imprinting stimulation-induced changes in the spatiotemporal patterns of evoked activity. After repetitive stimulation, the spatiotemporal evoked pattern during the late response becomes more similar to the pattern during the early evoked response. This modification lasted up to one hour after repeated stimulation. The increase of similarity between the late and early evoked responses after repeated sensory stimulation observed in anesthetized mice was also observed in wide-field extracellular glutamate recordings in head-fixed awake mice.

In summary, our results show that the changes induced by sensory experience to the component previously associated to sensory perception in primary sensory cortices might be strongly related to the late evoked responses. Since complex cognitive functions such as perception and learning involve the interaction of multiple brain structures, including subcortical regions, the study of experience dependent changes of spatiotemporal patterns of sensory evoked responses over the cortex can expand our understanding of such functions (Ferezou et al. 2007; Mohajerani et al. 2013; Luczak, McNaughton, et al. 2015; Karimi Abadchi et al. 2020). However, a more detailed study of the dynamics of cortical and subcortical structures activity might be needed to understand the brain mechanisms that are involved in perception and how they are modified by experience.

## Materials and Methods

### Animals

Twenty-seven C57Bl/6j adult (20-30 g, age 2-4month) mice were used for voltage-sensitive dye (VSD) experiments under anesthesia. For awake wide-field imaging experiments, 3 adult (>2 months) iGluSnFR transgenic mice (strain Emx-CaMKII-Ai85), expressing iGluSnFR in glutamatergic neocortical neurons (Marvin et al. 2013; Xie et al. 2016; Karimi Abadchi et al. 2020), were used. For this, Emx-CaMKII-Ai85 transgenic mice were generated by crossing the homozygous B6.129S2-Emx1tm1(cre)Krj/J strain (Jax no. 005628) and the B6.Cg-Tg(CamK2a-tTA)1Mmay/DboJ strain (Jax no.007004) with the hemizygous B6;129S-Igs7 tm85(teto-gltI/GDP*) Hze/J strain (Jax no.026260). This crossing is expected to produce expression of iGluSnFR within all excitatory neurons across all layers of the cortex, but not in GABAergic neurons (Huang and Zeng 2013; Madisen et al. 2015). Brain sections of the positive transgenic mice confirmed robust expression in the neocortex and hippocampus. All procedures were performed following approved protocols by the University of Lethbridge Animal Care Committee (ACC) and in accordance with the standards of the Canadian Council on Animal Care (CCAC).

### Surgery

#### Anesthetized experiments

Mice were anesthetized with 15% urethane in HEPES-buffered saline solution (1000-1250 mg/kg depending on the age and weight) and fixed in a stereotactic apparatus where the skull was rotated over the longitudinal axis of the skull 30 degrees to expose most of the dorsal and lateral cortex. Body temperature was increased to maintain 37°C with an electric heating pad regulated by a feedback thermistor throughout surgery and imaging. Mice were given Dexamethasone (80 μg) intramuscularly to prevent inflammation and Lidocaine (50 μl, at 0.2%) into the skin over the craniotomy area for local anesthesia. For Voltage-sensitive dye imaging experiments, a 7 × 6 mm unilateral craniotomy (bregma 2.5 to −4.5 mm, lateral 0 to 6 mm) was made and the dura mater was removed, as described previously (Mohajerani et al. 2010; Kyweriga et al. 2017; Bermudez-Contreras et al. 2018; Greenberg et al. 2018; Afrashteh et al. 2020). In all cases for VSD imaging, mice were also given a tracheotomy to assist with breathing. For each hour under anesthesia, the mouse was given an intraperitoneal injection of 10 ml/kg of 0.5% dextrose and 0.9% saline solution to maintain hydration.

#### Awake experiments

For recordings in awake mice, animals were anesthetized with isoflurane (2.5% induction, 1-1.5% maintenance), subcutaneous injections of 0.5gr/Kg buprenorphine half an hour before surgical procedures. A craniotomy of ~ 4mm of diameter was performed to expose the auditory cortex. This cranial window had the squamosal bone to its lateral end and its caudal end was 0.5mm anterior to the lambdoid structure. Finally, a stainless-steel head-plate was fixed to the skull using metabond and dental cement, a glass coverslip was placed on top to keep the surface clear from accumulating debris from the environment. Body temperature was increased to maintain 37°C with an electric heating pad regulated by a feedback thermistor throughout surgery. After two weeks of recovery from this procedure, these animals started to be habituated to the recording apparatus (see wide-field imaging procedure below).

### Wide-field optical imaging

For VSD imaging experiments, the dye RH-1691 (optical Imaging, New York, NY) was diluted in HEPES-buffered saline solution (0.5mg/1ml), applied to the brain for 45 min and rinsed subsequently, which stained all neocortical layers as reported previously (Mohajerani et al. 2010). The brain was then covered with agarose in HEPES-buffered saline at 0.6% concentration and sealed with a glass coverslip. This procedure reduced the movement artifacts produced by respiration and heartbeat. VSD imaging began ~30 min after washing unbound VSD. For VSD data collection, 12-bit images were captured at 150 Hz during evoked activity and at 100 Hz during spontaneous activity with a charge-coupled device (CCD) camera (1M60 Pantera, Dalsa, Waterloo, ON) and an EPIX E8 frame grabber with XCAP 3.8 imaging software (EPIX, Inc., Buffalo Grove, IL). The dye was excited using a red LED (Luxeon K2, 627 nm center) and excitation filters of 630 ± 15 nm. Images were taken through a macroscope composed of front-to-front video lenses (8.6 x 8.6 mm field of view, 67 μm per pixel). VSD fluorescence was filtered using a 673-to-703 nm bandpass optical filter (Semrock, New York, NY). To reduce potential VSD signal distortion caused by the presence of large cortical blood vessels, the focal plane was set to a depth of ~ 1mm from the cortex surface (Mohajerani et al. 2013). To monitor cortical activity in awake animals, the extracellular glutamate concentration was recorded in iGluSnFR mice, the same camera and lenses were used as for VSD recordings. However, a blue LED (Luxeon K2, 473 nm) and an excitation filter (Chroma, 467-499 nm) were used to excite the glutamate fluorescent indicators. The reflected fluorescent signal from excited indicators was filtered using a (Chroma, 510 to 550 nm) band-pass optical filter (Semrock, New York, NY). This sensor was used due to the high temporal resolution (similar to the VSD and better than calcium indicators) needed to capture the evoked activity dynamics in awake animals (VSD is not suitable to awake experiments). To reduce potential artifacts caused by the presence of large cortical blood vessels, the focal plane was set to a depth of ~1 mm from the cortical surface (Mohajerani et al. 2013). For awake glutamate recordings, mice were habituated to the recording setup after two weeks of recovery from the head-plate implant. This consisted of putting the animals one by one on the recording platform with one or two pieces of Cheerios cereal. After a few days of becoming familiar with the apparatus, the animals were head-restrained in incremental daily periods starting from 20 min, and increasing 5 minutes per day, reaching a total restriction time of 1.5 hours. During the head-fixation period, each animal was placed inside a plastic tube to limit motion and encourage relaxation. In addition, the temperature of the platform was increased to room temperature using microwavable heat pads.

### Sensory Stimulation

The experiment was divided into three different time periods. The first period consisted of 20 single-pulse stimulation trials of 5 seconds each. Each trial consisted of a 0.9 s baseline period followed by a single pulse stimulation, followed by 4.1 s of activity after stimulus onset. Single-pulse stimulation was given to three different sensory modalities in different experiments to evaluate changes in the evoked responses induced by repeated stimulation. To induce evoked responses in the somatosensory cortex (S1), a thin acupuncture needle was inserted into the paw and a 1ms of 0.2-0.3 mA electrical pulse was delivered. To induce visual evoked responses, a 1ms pulse of green light was delivered approx. 10 cm away from the contralateral eye of the recording hemisphere using a light emitting diode as described previously (Mohajerani et al. 2013). To induce auditory evoked responses, a 12 kHz 60-80dB 50ms tone was played to the contralateral side of the recorded hemisphere at 15-20 cm of distance. All auditory stimuli were generated with custom software in MATLAB (MathWorks, Natick, MA) at a sampling rate of 192 kHz using the RX-6 multi-function processor and output to an Electrostatic Speaker driver (Part# ED1, Tucker Davis Technologies, Alachua, FL), which in turn delivered the sounds to an electrostatic speaker (Part# ED-10, Tucker Davis Technologies, Alachua, FL). The speaker was placed 10 cm directly to the left side of the animal’s head (contralateral of the recorded hemisphere). The second period consisted of repeated continuous somatosensory or auditory stimulation for 30 min. For repeated somatosensory stimulation, electrical pulses were given to the hindpaw as a continuous 20 Hz frequency stimulation train of pulses (0.2-0.3 mA, 1ms). For repeated auditory stimulation, a series of 12 kHz 1 sec tone followed by 1 sec of silence were given for 30 min. Finally, the third period consisted of a set of 20 single-pulse stimulation trials of 5 seconds each. Each trial was the same as in the first period.

### Brain state manipulation

Before brain imaging started, Methamphetamine (1mg/kg, Sigma-Aldrich) was injected to induce a desynchronized brain state to modify the patterns of cortical activity of the sensory evoked responses by repeated stimulation. In previous work, we have demonstrated that even in anesthetized animals, it is possible to induce stimulus dependent modifications to the patterns of cortical activity by repeated stimulation of the periphery when the cortex is in a desynchronized state (Bermudez Contreras, Gomez Palacio Schjetnan, et al. 2013). To ensure that the brain reached a desynchronized brain state, data collection started 10-15 min after injection for the drug effect to stabilize and we visually verified that the cortical LFP showed increased content in high frequency and reduced amplitude. It is known that desynchronized brain states enhance changes induced by repeated stimulation during urethane anesthesia.

### Data analysis of imaging data

To correct for time-dependent changes in VSD signals due to bleaching, 20 non-stimulation interleaved trials were used for normalization of the evoked data. A 10 s interval between each sensory stimulation trial was used. Although VSD fluorescence has been shown to have relatively high labeling at a depth of ~750 μm across the cortex (Mohajerani et al. 2010), all VSD recordings were expressed as a percentage change relative to baseline VSD responses (ΔF/F0 × 100%) to reduce regional bias in VSD signal caused by uneven dye loading or brain curvature where ΔF = (F − F0), F is the fluorescence signal at any given time and F0 is the average of fluorescence over baseline frames. Analogously, for glutamate imaging changes in glutamate concentration were also defined as a relative quantity to a baseline (ΔF/F0 × 100%).VSD imaging of spontaneous activity was continuously recorded in the absence of sensory stimulation at 100 frames per second. Slow, time-dependent reductions in VSD fluorescence were corrected in MATLAB® using a zero-phase lag Chebyshev bandpass filter (zero-phase filter) at 0.1 to 6 Hz. Ambient light resulting from VSD excitation (630 nm) was measured at 8.65 × 10−3 W/m2. In addition, for glutamate awake recordings, raw data were corrected using global signal regression to remove global hemodynamic and illumination fluctuations (Chan et al. 2015; Xie et al. 2016).

Sensory stimulation was used to determine the regions of interest (ROIs) for the primary sensory areas (HLS1, FLS1, V1 and A1), and secondary somatosensory areas (HLS2, FLS2). All the wide-field imaging recordings were registered to the Allen Brain atlas coordinates matching the 30 degrees rotation over the longitudinal axis of the skull and using the initial sensory evoked responses ROIs as reference points to perform 2D mapping transformation using the *fitgeotrans* function in MATLAB. To obtain the dynamics of the evoked responses, the imaging signal was averaged over a 15 pixel-diameter circle centered in the initial activation area for each recording (Figs. 1B, 2B, 3B and 4B). Similar results were obtained with different ROI sizes and having independent ROIs for the before and after conditions.

**Figure 1.**
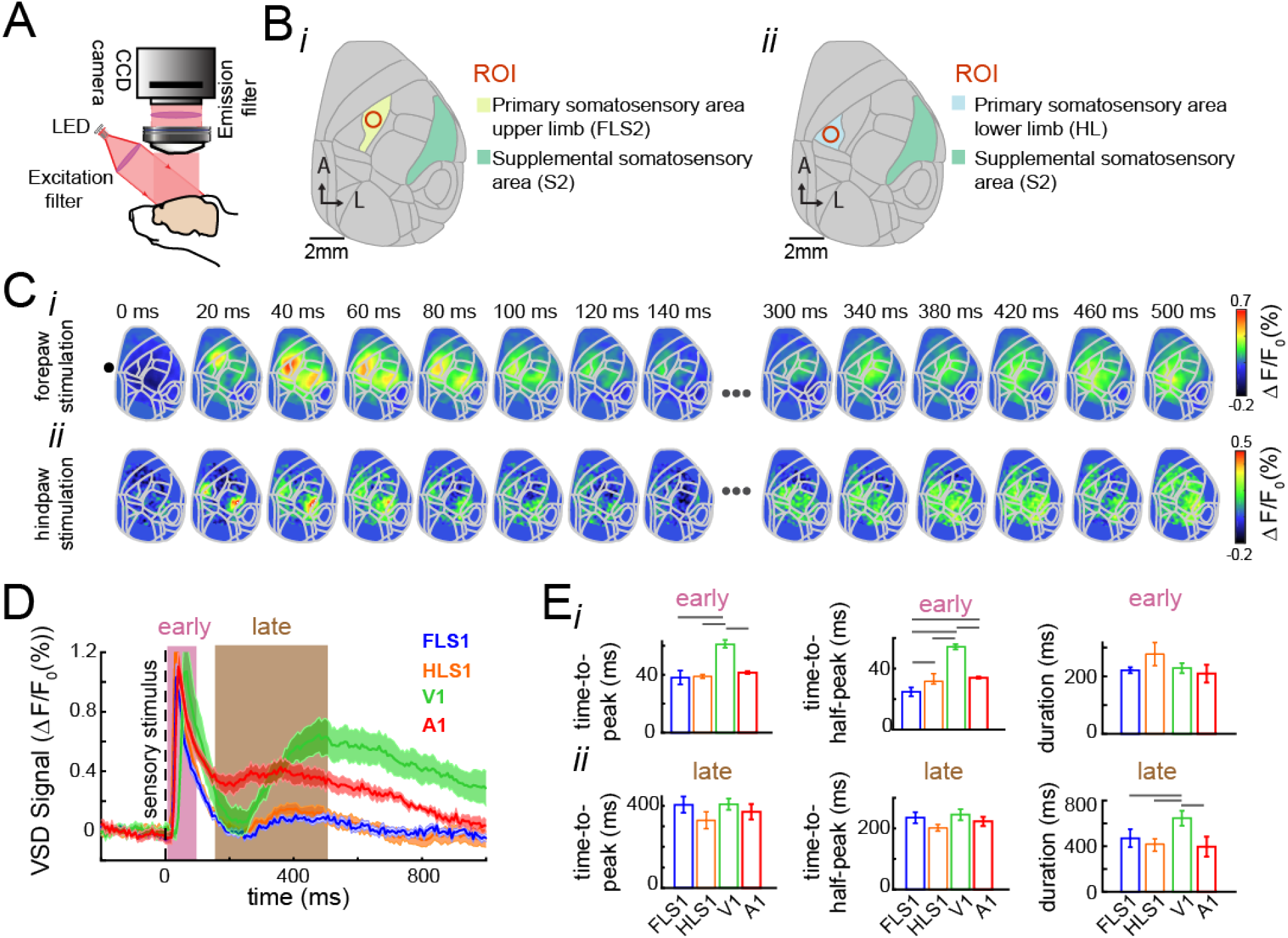
Spatiotemporal organization of the early and late sensory evoked responses in mouse cortex. **(A)** Schematic of the VSD imaging apparatus. A CCD camera records changes in voltage over the imaged area. The voltage-sensitive dye is excited by a red LED, the reflected light is then filtered and finally captured by the camera. **(B)** Cortical maps registered to the Allen Mouse Brain atlas. Region of interests (ROIs) for forepaw (i) and hindpaw stimulation (ii) correspondingly. **(C)** Example of the spatiotemporal evoked pattern to forepaw stimulation (i) and hindpaw stimulation (ii). **(D)** Average evoked responses to stimulation of the forepaw (FL), hindpaw (HL), visual stimulation (VC) and auditory stimulation (AC) before repeated stimulation. Solid lines represent the mean (n=7, 12, 7, 12 for each sensory modality respectively). The shadows represent the SEM across animals. FLS1, HLS1 denote responses to forelimb and hindlimb stimulation recorded in the forelimb and hindlimb areas of primary somatosensory; V1 denotes responses to visual stimulation recorded in the primary visual cortex; A1 denotes the response to tone stimulation recorded in primary auditory cortex. **(E)** Temporal organization of the evoked response for different sensory modalities as in D. Time-to-peak, time-half-to-peak, and duration for the early (top row) and late (bottom row) evoked responses. The error bars denote the SEM. The gray horizontal lines denote significant differences across the populations for the different stimulated peripheries (p < 0.05, one-way ANOVA with Bonferroni correction, n=9, 12, 8, 10, respectively).

**Figure 2.**
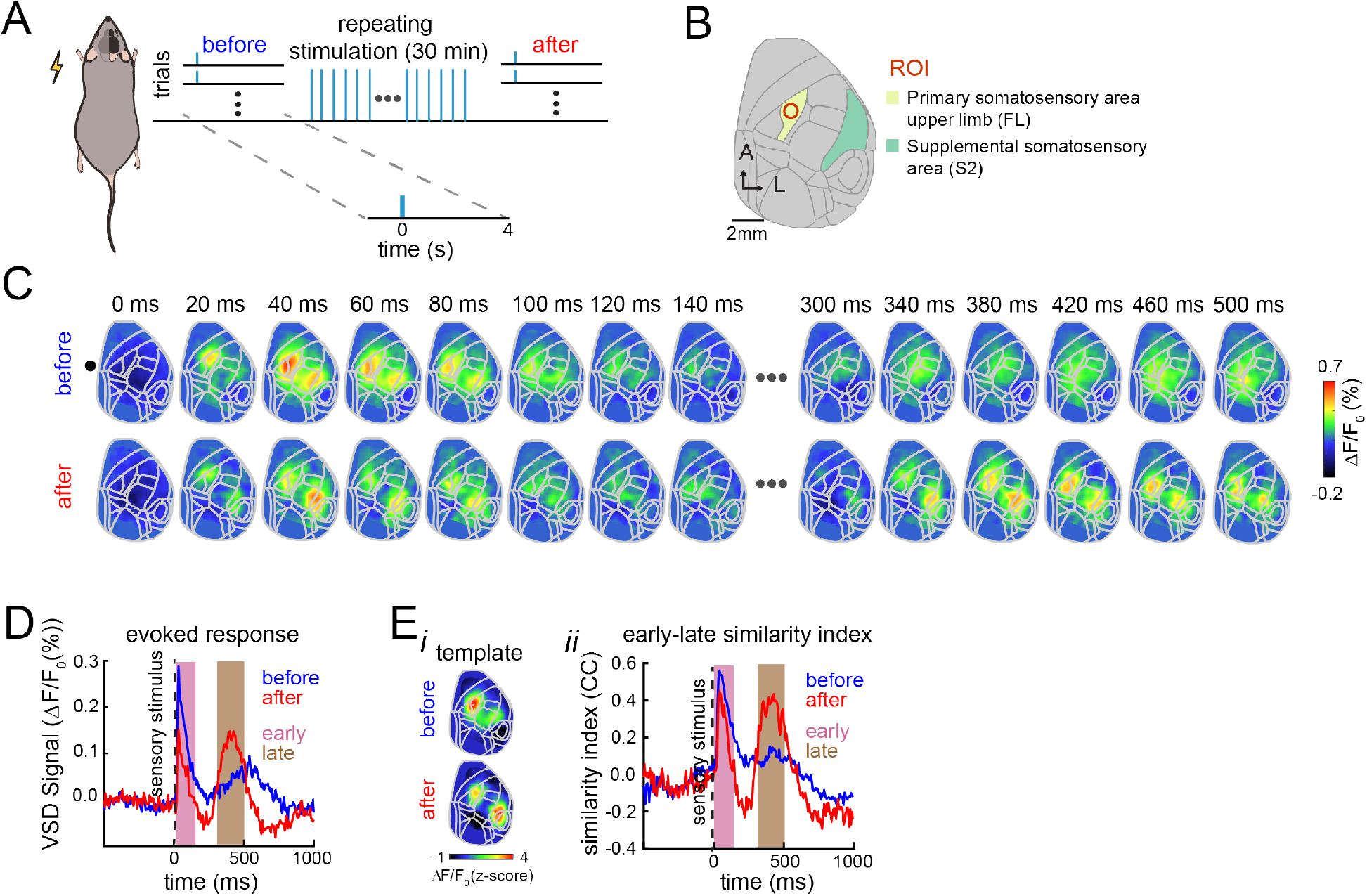
Spatiotemporal changes in the early and late evoked responses induced by repeated stimulation. **(A)** Experimental protocol. The anesthetized mice injected with amphetamine received interleaved single-pulse electrical stimulation to the forepaw and a LED flash as visual stimulation (20 trials each) followed by 30 min of repeated stimulation of the forepaw at 20 Hz and finally received another interleaved single-pulse forepaw and visual stimulation (20 trials each). **(B)** Cortical map registered to the Allen Mouse Brain Atlas for our wide-field recordings. The yellow region corresponds to the forelimb area of the primary somatosensory area and the green region corresponds to the secondary somatosensory area. **(C)** Example of trial-averaged evoked activity pattern in response to electrical fore paw stimulation (i) before (top row) and after (bottom row) repeated forepaw stimulation. **(D)** Average response in the FLS1 area (ROI) to forelimb electrical stimulation before (blue) and after (red) repeated stimulation across trials (n=30 trials, one animal). **(E)** Trial-average template similarity of evoked responses for the same mouse. (i) Templates of early evoked activity before and after repeated stimulation. (ii) Trial-average similarity between the templates and the evoked response before (blue) and after (red) repeated stimulation.

**Figure 3.**
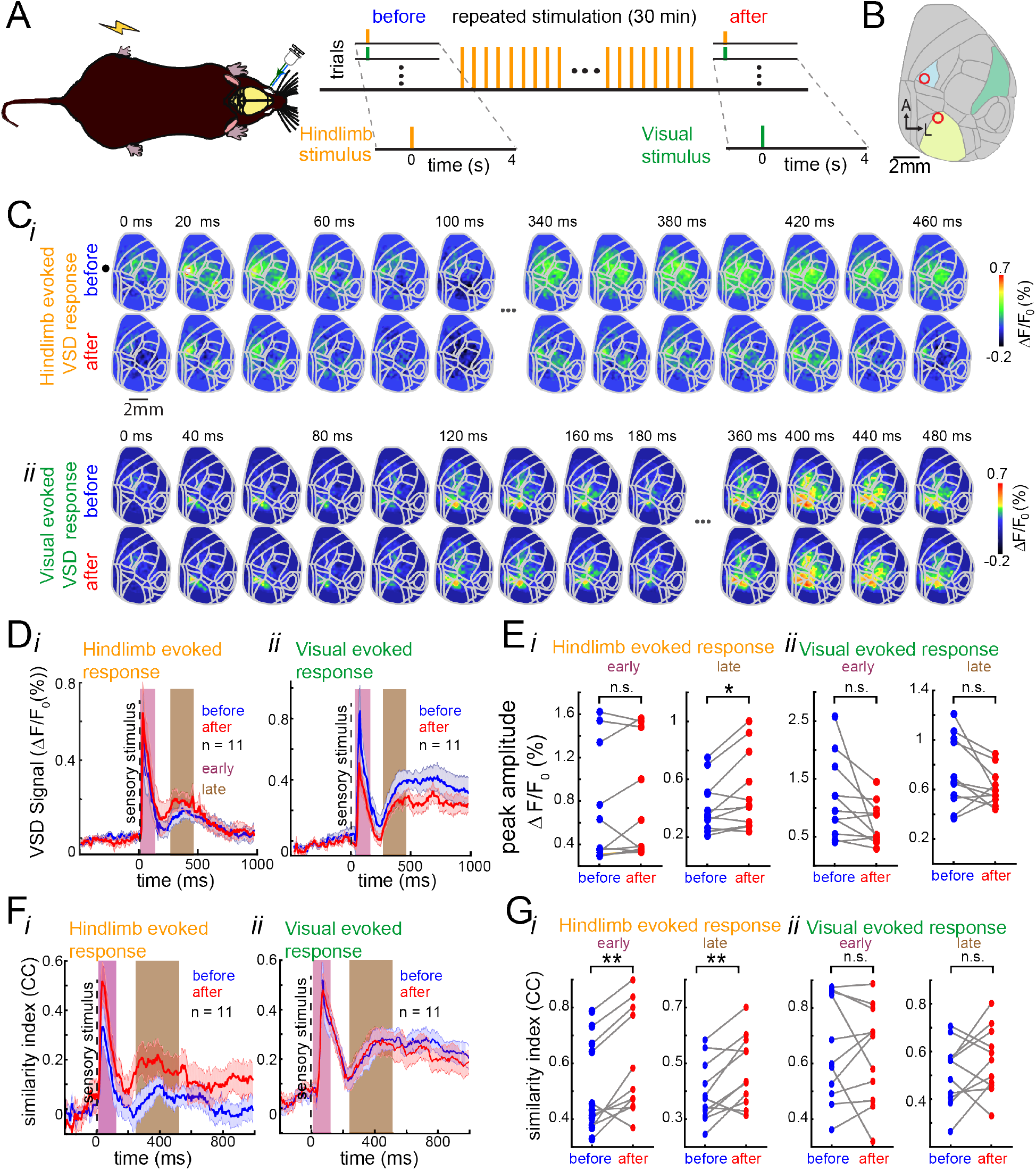
Spatiotemporal changes in the early and late evoked responses are stimulation dependent. **(A)** Experimental protocol. The anesthetized mice injected with amphetamine received interleaved single-pulse electrical stimulation to the hindpaw and a LED flash as visual stimulation (20 trials each) followed by 20 min of repeated stimulation of the hindpaw at 20 Hz and finally received another interleaved single-pulse hindpaw and visual stimulation (20 trials each). (B) Cortical map registered to the Allen Mouse Brain Atlas for our wide-field recordings. The blue region corresponds to the hindlimb area of the primary somatosensory area, the yellow region corresponds to the primary visual area. **(C)** Example of trial-average evoked activity in response to hindlimb pulse stimulation (i) and visual stimulation (ii) before (top) and after (bottom) hindpaw repeated stimulation in the same mouse. **(D)** Average evoked VSD response across animals before (blue) and after (red) hindpaw repeated stimulation in response to hindpaw stimulation (i) and to visual stimulation (ii). The shaded regions denote the S.E.M. (animals n=11). Green and orange regions denote the early and late components of the evoked responses, respectively. **(E)** Paired comparison of the peak amplitude of the early and late evoked response before and after repeated stimulation for hindpaw stimulation (i) and visual stimulation (ii) for all animals (n=11). **(F)** Average template similarity between early (left) and late (right) evoked responses to pulse hindpaw stimulus (i) and visual stimulus (ii) before (blue) and after (red) repeated hindpaw stimulation across animals. **(G)** Paired comparison of the template similarity of early and late evoked responses to hindpaw (i) and visual stimulation (ii) before (blue) and after (red) during early (left) and late components (right) for all animals. * and ** represent significant increase in amplitude of the late evoked response after repeated hindpaw stimulation (t-test, p < 0.05 and p < 0.01 respectively).

**Figure 4.**
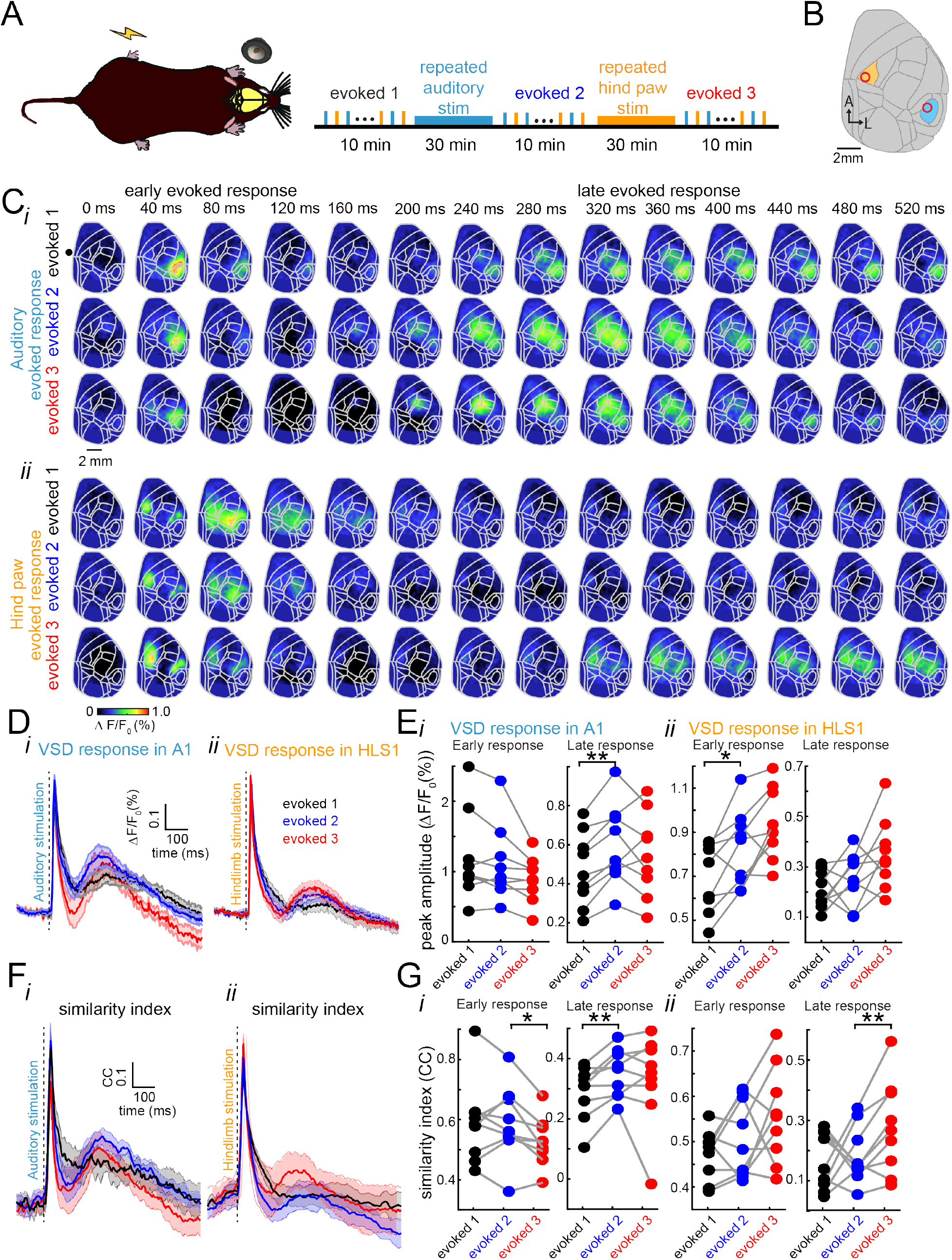
Spatiotemporal changes in the early and late evoked responses induced by repeated stimulation of different sensory modalities. **(A)** Schematic of the experimental protocol. The anesthetized mice injected with amphetamine received both repeated auditory and repeated hindpaw stimulation. **(B)** Auditory and hindpaw repeated stimulation in the same preparation. Single-pulse evoked responses to auditory and hindpaw stimulation were interleaved every 10 sec before any repeated stimulation (evoked1), after auditory repeated stimulation (evoked2) and after hindpaw repeated stimulation (evoked3). **(C)** Trial-average evoked responses to auditory stimulation (i) and hindpaw stimulation (ii) before (evoked1) and after (evoked2) repeated auditory stimulation and after repeated hindpaw stimulation (evoked3) for one mouse. **(D)** Average evoked response in the auditory cortex (i) and in the hind limb area of the somatosensory cortex (ii) across animals (n=9 animals). **(E)** Pair-wise comparison between the peak amplitude of the responses to auditory stimulation (i) and hindpaw stimulation (ii) during the early (left) and late (right) components for all animals. **(F)** Mean similarity index for auditory early and late evoked responses (i) and hindpaw evoked responses (ii) before any repeated stimulation (black), after repeated auditory stimulation (blue) and after repeated hindpaw stimulation (red) across animals (n=9). **(G)** Paired comparison between the similarity between the early and late auditory (i) and hindpaw (ii) evoked responses during the early (left) and late hindpaw evoked responses (right) before any repeated stimulation (black), after auditory repeated stimulation (blue) and after repeated hindpaw stimulation (red). (n=9, each line denotes an animal. * and ** denote p<0.05 and p<0.01, paired t-test respectively).

To measure the similarity between the early and late evoked responses, each stimulation trial was compared against a template constructed from the average evoked response during the first 33.3 ms (5 frames) after the time when the amplitude of the evoked response was 2 STD over the baseline. Similar results were obtained when using different templates lengths (data not shown). The similarity between the template and the evoked activity was calculated as the 1D correlation coefficient between the template and each frame for each stimulation trial. To avoid the similarity being influenced by the amplitude of the evoked responses, we applied a z-score transformation to the evoked activity (and template) before calculating the correlation coefficient. To measure the similarity of the spontaneous activity to the early evoked response, the same template matching procedure was used. The similarity between the spontaneous activity and the template was calculated using a sliding window over 1.5 sec of the inter-stimulus interval after 2.5 sec of the stimulus onset of each trial (Fig. 5A-B).

**Figure 5.**
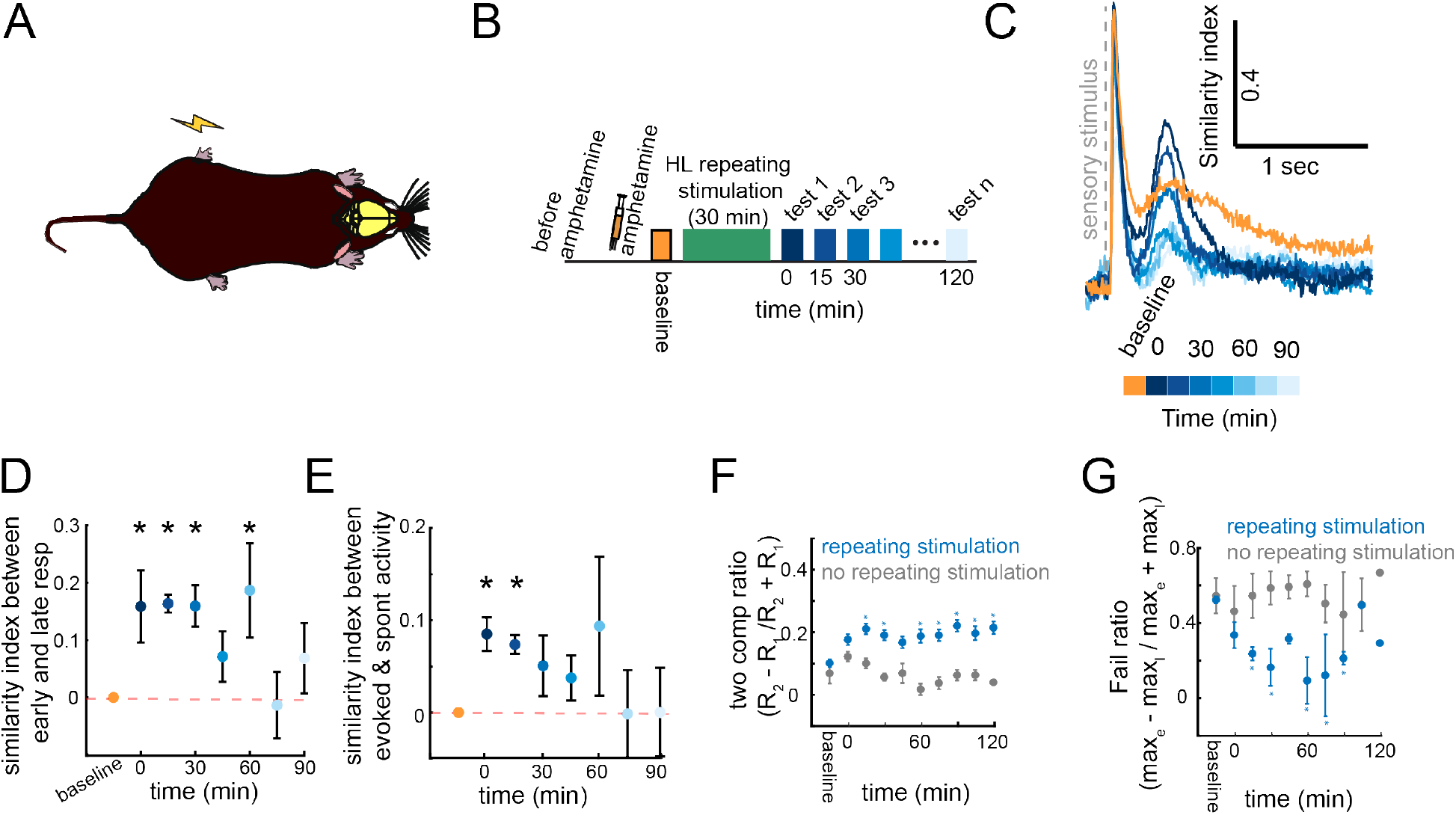
Dynamics of changes induced by repeated stimulation. **(A)** Experimental setup. Anesthetized animals received single pulse electrical stimulation to the hindpaw before repeated stimulation (baseline) and after. **(B)** Sensory evoked responses were tested every 15 min for 1.5 hr after repeated stimulation. **(C)** Average similarity index between early and late spatiotemporal evoked responses in a single animal (n=30 trials)**. (D)** Average similarity between the early and late components of evoked responses for multiple mice (n=8 animals, error bars represent S.E.M. and stars represent statistically different to baseline trials, one-way ANOVA with Bonferroni correction). **(E)** Average similarity of evoked patterns of activity during spontaneous activity measured up to 1.5 hours after repeated stimulation across animals (same as f). **(F)** Dynamics of the two-component ratio using a mixture of Gaussians model. Error bars denote the S.E.M. and stars denote trials that are significantly different to the baseline trials (ANOVA with Bonferroni correction, n= 90 trials). **(G)** Dynamics of the Fail ratio between the early and late component for animals that received repeated stimulation (blue) and animals that did not receive repeated stimulation (grey).

To evaluate if the amplitude of the early and late components were related, the Fail ratio was defined as the division between the difference of the maximum amplitude of the early response and the maximum amplitude of the late response, divided by the sum of the maximum amplitude of the early response and the maximum amplitude of the late response.

### Statistical tests

One-sided paired t-tests and Wilcoxon signed-rank tests (when distributions were not normal) were used to determine if the similarity between the early and late evoked responses (in the same animal) increased after repeated stimulation across all animals. When more than two groups were compared, ANOVA with ad-hoc Bonferroni correction was used. All statistical tests were performed using MATLAB built-in functions.

### Code and Data availability

All code was developed in MATLAB (MathWorks) and is available on github.com. Data will be made available upon request.

## Results

### Sensory evoked responses are organized into early and late components

Brain activity was monitored using VSD wide-field imaging over much of the dorsal cortex of the right hemisphere in anesthetized mice injected with amphetamine (Figs. 1A-B). Consistent with previous wide-field imaging studies, we found that stimulation of the fore and hind paws resulted in evoked responses first in the forelimb and hindlimb area of the primary somatosensory cortex (S1), followed by an activation of the secondary somatosensory area (S2) (Fig. 1C) (Lim et al. 2012; Mohajerani et al. 2013). Moreover, we found that the sensory evoked response to a single pulse stimulation consists of two components, an early and a late evoked responses, as reported in other studies (Sachidhanandam et al. 2013; Funayama et al. 2015a; Manita et al. 2015a; Bermudez-Contreras et al. 2018). This bimodal organization is observed across different modalities (Fig. 1D) but with different temporal characteristics (Fig. 1E), particularly for visually evoked responses. Similar dynamics have been reported previously (Mohajerani et al. 2013; Xie et al. 2016). For example, the early evoked response lasts 240 ± 7.7 ms on average for hindpaw electrical stimulation and 38.8 ± 1.3 ms time-to-peak from stimulus onset (n=9 animals). The source of the early evoked response is attributed to thalamo-cortical connections (Manita et al. 2015a). The late component of the hindpaw evoked response consists of a larger period of time that on average lasts 410.3 ± 28.5 ms and with a time-to-peak of 318 ± 68 ms from stimulus onset (n=9 animals).

### Sensory evoked responses are modified by repeated sensory stimulation

To evaluate the changes induced by sensory experience, the evoked responses to single pulses of stimulation were recorded before and after repeated intermittent electrical stimulation of the forepaw (Fig. 2A-B). After 30 min of repeated stimulation of the forepaw, a reorganization of the evoked response to single pulse stimulation was observed (Fig. 2C). The late evoked response increases in amplitude after repeated stimulation and the early component tends to decrease (Fig. 2D). In addition, after repeated stimulation, the late evoked pattern of activity resembles the early evoked response more closely than before (Fig. 2C). To quantify this spatiotemporal reorganization of the late evoked response, we calculated the similarity between the early and late evoked responses for each stimulation trial using template matching. The similarity was calculated as the correlation coefficient between a template from the average response during the first 33.3 ms and each stimulation trial (Fig. 2E i-ii). After repeated stimulation of the forepaw, the similarity between the early and late components increases.

### Stimulation dependent changes in the amplitude of the early and late somatosensory evoked responses

To evaluate whether the changes induced by stimulation occur only in the sensory modality that received the repeated stimulation (i.e. input selectivity), electrical pulses to the hindpaw of the animal were interleaved with visual stimulation (see Methods), and repeated electrical pulses of stimulation (Fig. 3A-B) were only given to the hindpaw (not repeated visual stimulation).

After repeated electrical stimulation of the hindpaw, an overall increase in the average amplitude evoked response (across animals) was observed, mainly during the late component of the hindpaw evoked response (Fig. 3C (i), 3D (i) and 3E (i)). Interestingly, the opposite effect is observed in the visual evoked response. That is, the amplitude of the late visual evoked response tends to decrease after repeated hindpaw stimulation. However, this comparison did not yield a significant change (Fig. 3C (ii), 3D (ii) and 3E (ii)). The quantification of the amplitude of the average evoked response shows that there is a significant increase (pair-wise t-test, p <0.05, n=11) in the amplitude of the late hindpaw evoked response after repeated stimulation (Fig. 3E (i)) but no significant changes in the visual evoked response (Fig. 3E (ii)).

Although we tried to provide repeated visual stimulation in this preparation, we could not find an adequate visual stimulation protocol that would induce changes in the visual evoked responses that we could capture following the methodology described for somatosensory and auditory sensory modalities. It is important to note that the characteristics of visually evoked cortical responses seem different to the other sensory modalities (Fig. 1D-E). More experimental exploration of such protocols (duration of stimulus and frequency) might be necessary to achieve similar results to the ones from the hind and fore paws and auditory stimulation.

### Reorganization of the late somatosensory evoked response after repeated stimulation

To evaluate whether the reorganization of the spatiotemporal late evoked response after repeated stimulation was exclusive to the sensory modality that was stimulated, the similarity between the late evoked response and the pattern of activity during the early response were compared for every trial of hindpaw and visual stimulation.

The similarity between the early and late evoked responses was calculated using template matching as explained previously. After hindpaw repeated stimulation, an increase in the mean similarity across all animals between the early and late hindpaw evoked responses was observed (Fig. 3F (i)) but not in the visually-evoked responses (Fig. 3F (ii)). Moreover, there is a significant increase in the mean similarity between the early and late evoked responses across trials after repeated hindpaw stimulation but not for the visual evoked responses (Fig. 3G, paired t-test, p < 0.05, n=11).

In summary, these results suggest that the spatiotemporal reorganization of the spatiotemporal evoked patterns after repeated sensory stimulation is input-selective and occurs in the corresponding evoked activity pattern.

### Spatiotemporal reorganization of the late evoked response in two sensory modalities

To evaluate whether the changes induced by the repeated stimulation were restricted to the somatosensory cortex or could be observed in other sensory cortices, auditory and hind paw repeated stimulation were provided in the same preparation (Fig. 4A-B). As before, the single-pulse evoked responses were compared before and after repeated sensory stimulation. This time, a 50 ms 12 Khz tone was interleaved with 1 ms 300 μA pulse to the hind paw, separated by 10 seconds. These single pulse stimulations (20 trials each) were repeated three times: one before any repeated stimulation (denoted evoked1), one after repeated auditory stimulation (denoted evoked2) and one more after repeated hind paw stimulation (denoted evoked3) (Fig. 4A). The spatiotemporal patterns of activity show qualitatively, that after single tone stimulation, the early response activity starts in the auditory cortex after 20 ms from stimulus onset and expands for approximately 120 ms. Similarly, the late auditory evoked response starts around 200 ms after the stimulus onset and lasts for 300 ms approximately (Fig. 4C (i)). Analogously, the hind paw early evoked response starts after approximately 20 ms after stimulus onset in the primary somatosensory cortex and lasts for 100 ms including the secondary somatosensory cortex and expanding to the midline areas (Fig. 4C (ii)). Quantitatively, our analyses indicate that before repeated stimulation, the early auditory evoked response lasted on average 214 ms ±9.6 SEM and 240 ms ±7.7 SEM for hindpaw stimulation. After repeated tone stimulation, the early auditory and early S1HL evoked response lasted on average 226.6 ms ±13.7 SEM and 200 ms ±5.4 SEM respectively (n=9 animals). Before repeated stimulation, the late auditory and S1HL evoked responses lasted on average 454 ms ±29.8 SEM and 510.3 ms ±28.5 SEM for hindpaw stimulation respectively. After repeated stimulation, the late tone and hindpaw evoked responses lasted 518.2 ms ±24.5 SEM and 645.8 ms ±24.4 SEM respectively (Fig. 1 D-E, Fig. S1 B).

We found that after repeated auditory stimulation the amplitude of the early auditory evoked responses does not change compared to baseline (Fig. 4D (i) and 4E (i), left). However, the amplitude of the late auditory evoked responses increased after repeated auditory stimulation (Fig. 4D (i), and 4E (i), right). In contrast, the amplitude of the late hindpaw evoked responses did not increase after the auditory repeated stimulation (Figs. 4D (ii), 4E (ii) right). After repeated hindpaw stimulation (evoked3), the early and late amplitude of the auditory evoked response (red trace) tends to decrease, although these changes were not statistically significant (Fig. 4D (i), 4E (i)). In contrast, the early and late hindpaw evoked response increased (Fig. 4D (ii)), but these changes were not statistically significant (Fig. 4E (ii)). We were expecting to observe an increase of the amplitude of the late auditory evoked response after repeated stimulation, this lack of an increase in amplitude might be due to a ceiling effect. More experiments are needed to explore this possibility.

To evaluate whether the similarity between the early and late spatiotemporal responses was modified after repeated sensory stimulation, the evoked responses were compared against a template built from the first 33.3 ms after the onset of the evoked response. As previously, we calculated the correlation coefficient between the spatiotemporal auditory evoked response for every trial and the template of the early auditory evoked response before (evoked1, black trace) and after auditory repeated stimulation (evoked2, blue trace). Analogously, we compared the similarity between the early and late hindpaw evoked responses before and after repeated hindpaw stimulation (evoked3, red trace) (Fig. 4F-G). A significant increase in the similarity between the auditory late and early components of the evoked response was observed after repeated auditory stimulation but not after repeated hindpaw stimulation (Fig. 4G (i)). Analogously, a significant increase in the similarity between the late and early components of the hindpaw evoked response was observed only after repeated hindpaw stimulation (Fig. 4G (ii)) but not in the auditory evoked responses. In summary, these results show that repeated sensory stimulation causes a reversible reorganization of the evoked responses. This reorganization consists of an increase in the similarity between the spatiotemporal late evoked response and the early evoked pattern of activity. Moreover, this reorganization of the cortical activity occurs in more than one sensory modality at the mesoscale level.

Altogether, our results show that the evoked response is formed by an early and a late component at the mesoscale level. After repeated sensory stimulation, the similarity of the spatiotemporal pattern of the early and late components of the evoked response increases. Moreover, our results indicate that this increase in similarity is due to the changes of spatiotemporal pattern of the late component to closely resemble the pattern of activity during the early evoked response. In addition, these changes are exclusive to the sensory modality that receive repeated sensory stimulation and, finally, these changes occur selectively in different sensory modalities.

### Duration of the cortical activity reorganization induced by repeated sensory stimulation

To evaluate how long these changes last for, the evoked responses to single pulse stimulation of the hindpaw were recorded up to 90 min after repeated stimulation (Fig. 5A-B). The similarity between the early and late evoked responses last for up to one hour after repeated stimulation compared to their similarity before (baseline) repeated stimulation (Fig. 5C-D) (one-way ANOVA with Bonferroni correction, the stars represent p<0.05). To examine whether the stimulation affected subsequent spontaneous activity, we calculated the spatial correlation between the sensory-evoked VSD signals (templates) and spontaneous VSD signals. We found that repeated hindpaw stimulation can have lingering effects on the regional patterns of spontaneous activity (after stimulation ceases). Our analysis indicate that evoked-like activity reverberates during the subsequent periods of spontaneous activity up to 15-30 min after the repeated stimulation stopped (Fig. 5E). It is important to note that the comparison was carried out against the baseline similarity, which corresponds to spontaneous activity periods before repeated stimulation. Moreover, the organization of the evoked response in two components (e.g. increase in amplitude of the late evoked response) is observed even 90min after repeated stimulation. These changes in organization of the evoked responses are not observed in animals that did not receive repeated stimulation (Fig. 5F, G).

### Sensory experience dependent changes of auditory evoked responses in awake mice

To evaluate whether the characteristics of the evoked responses changed under anesthesia, we compared the evoked responses in the same animals when they were awake and under urethane anesthesia. In this experiment, we monitored the wide-field extracellular changes in glutamate using Emx-CaMKII-Ai85 strain mice (Xie et al. 2016; Karimi Abadchi et al. 2020). We found that the temporal organization of the evoked responses remained the same when comparing the correlation coefficients of t-Distributed Stochastic Neighbor Embedding (t-sne) projection or different pairs of principal components using Principal Component Analysis (PCA) of the evoked traces (Fig. S2 A-C) (using Fisher’s z transformation (Fisher 1925)). However, the evoked responses were faster when the animals were awake (Fig. S2 D). It is important to note that previous studies have reported no significant differences between the peak amplitude, the time to peak and decay time of the evoked responses in A1 or V1 between awake and lightly-anesthetized mice using VSD (Mohajerani et al. 2013). In addition, it is also important to mention that, in contrast with VSD imaging, the dynamics of the late evoked responses in our glutamate recordings in iGluSnFr mice might be contaminated with changes in blood oxygenation induced by the sensory stimulation (Grinvald and Hildesheim 2004; Xie et al. 2016). This might be an issue as the blue excitation light in these recordings might interfere with the green emission fluorescence by changes in hemoglobin (in contrast with the red-shifted wave length excitation of VSD) that might cast doubts of the organization of the evoked responses in awake animals (Xie et al. 2016). However, other electrophysiological studies in awake mice in which brain activity is not contaminated with changes in blood oxygenation, report the same temporal organization of the evoked responses (Funayama et al. 2015b) and confirmed by others (Bermudez Contreras, Gomez Palacio Schjetnan, et al. 2013; Bermudez-Contreras et al. 2018).

To evaluate whether the changes in evoked responses caused by repeated sensory stimulation were exclusive to anesthetized mice, we also monitored the wide-field extracellular changes in glutamate in awake head-fixed transgenic Emx-CaMKII-Ai85 mice before and after repeated tone stimulation (Fig. 6A-B). Similarly to the anesthetized experiments, repeated auditory (tones) stimulation induced a modification of the spatiotemporal patterns of cortical activity in awake animals, in this case, in the auditory cortex (Fig. 6 C). We quantify how these patterns were modified by repeated stimulation, the amplitude of the evoked responses and the similarity between the early and late components of the evoked response were compared before and after repeated auditory stimulation as in the anesthetized experiments. In contrast to the anesthetized experiments, the peak amplitude of the auditory evoked responses (not early not late) was not significantly modified after repeated auditory stimulation (Fig. 6D). However, there is an increase of the similarity between the template (similarity index) only in the late evoked response after repeated stimulation (Fig. 6E).

**Figure 6.**
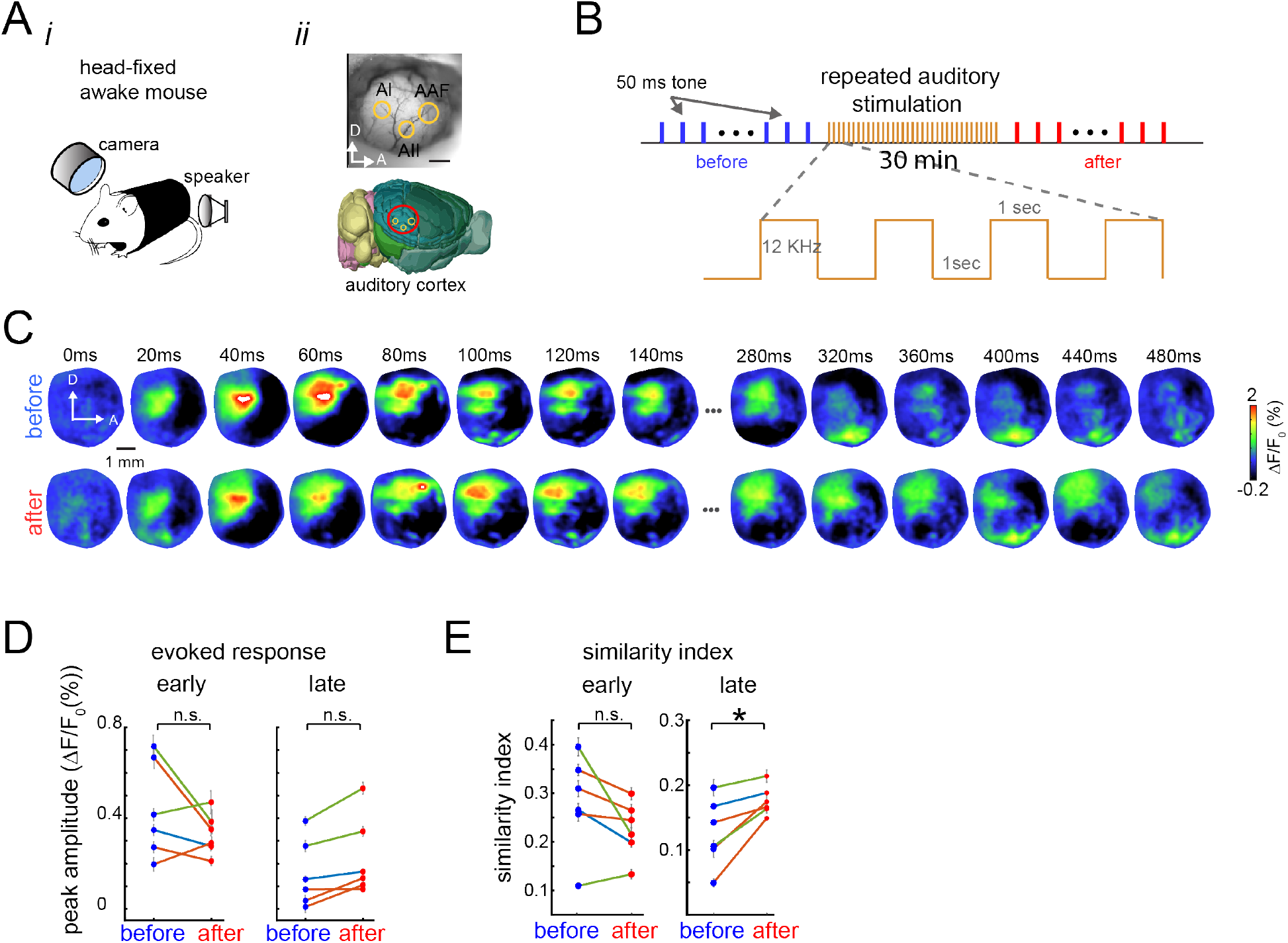
Sensory experience-dependent changes to the patterns of evoked activity in awake mice. **(A)** Experimental setup. Animals are head-fixed passively listening to tones (i) while brain activity is recorded in the auditory cortex (ii). **(B)** Experimental protocol. Single 50 ms tone trials were recorded before (blue) and after 30 min of repeated auditory stimulation (red) **(C)** Representative spatiotemporal pattern of trial-average evoked activity before (top row) and after (bottom row) repeated auditory stimulation for one mouse. **(D)** Mean peak amplitude of the auditory evoked response during the early evoked response (left) and late evoked response (right), before (blue dots) and after (red dots) repeated auditory stimulation (colored lines represent sessions for 3 different animals. **(E)** Similarity between the sensory-evoked template and early evoked responses (left) and late evoked responses (right) in awake animals before (blue) and after (red) repeated stimulation. Each session had 30 trials (n=30) of auditory stimulation Error bars denote S.E.M. and stars denote p < 0.05, paired t-test).

## Discussion

The sensory evoked response is formed by two components. The first component consists of the initial evoked response that, depending on the sensory modality, depending on the sensory modality, lasts 200 ms on average after stimulus onset (early evoked response). The second component consists of an additional ‘bump’ of activity that starts around 200 ms after stimulus onset and lasts from 200 ms up to 600 ms depending on the sensory modality (Fig. 1D-E). A similar description of the evoked response in several sensory areas has been reported (Sachidhanandam et al. 2013; Crochet and Petersen 2015; Funayama et al. 2015a; Manita et al. 2015a). In these studies, it is argued that the second component of the evoked response is associated with sensory perception.

In this work we demonstrate how repeated stimulation alters the dynamics of the evoked responses to sensory stimulation at the mesoscale level. We found that after repeated stimulation, the amplitude of the late evoked response increases after repeated sensory stimulation and the spatiotemporal pattern of the late evoked response becomes more similar to the pattern of the early evoked response. These experience dependent modifications of spatiotemporal brain activity occur in the sensory modality that was stimulated (i.e. somatosensory and auditory cortices) and can be induced in more than one modality in the same preparation and, in both, anesthetized and awake animals. In summary, we show that most of the changes induced by the repeated stimulation occur during the late component of the evoked responses and these changes make the patterns of activity of the late component more similar to the pattern of activity of the early evoked response.

### The bimodal organization of the sensory evoked response

According to the literature, the two components of the sensory evoked response are produced by different brain circuits and are related to different functions. On the one hand, the early component is thought to be produced by the feedforward thalamocortical inputs to the corresponding sensory areas (Manita et al. 2015a) and is associated with sensory stimulus identity. Another possibility is that they could be originated by direct inputs from brainstem or even spinal cord (Kiritani et al. 2012). The second component is thought to be produced by a combination of feedback cortico-cortical and thalamocortical inputs (Guillery and Sherman 2002; Sachidhanandam et al. 2013; Manita et al. 2015a) or by the cortico-thalamic-striatal loop (Mandelbaum et al. 2019) and it is associated with sensory perception. At least for the cortical feedback, it has been suggested that the late evoked response in primary sensory areas is originated by a neuronal population in higher cortical areas and that this signal is associated to perceptual processing (Kwon et al. 2016; Yamashita and Petersen 2016; Romo and Rossi-Pool 2020).

Apart from the perceptual content of the biphasic organization of cortical evoked responses, there are other ideas regarding the function of the biphasic organization of cortical evoked responses. Recently, a similar biphasic organization of cortical activity has been proposed as an information organization in which brain activity is transmitted in packets. The early component is proposed to contain categorical (more general) information and be less variable. The late component is proposed to contain more specific information (identity), and therefore is more stimulus specific (Luczak, Mcnaughton, et al. 2015).

Another possible reason for the biphasic evoked response could be provided from the neurophysiology of top-down modulation of evoked responses in primary sensory areas (Friston 2018). Similar modulation occurs in reinforcement learning mechanisms in the brain (Roelfsema and Holtmaat 2018). In this view, during learning, a feedback signal is necessary to perform the synaptic updates (strengthened or weakened) of the circuits according to rewards or punishment associated with a selected action triggered by the received stimulus. Although, in our experiments, there was not explicit reward, the late component of the evoked response, arguably a signal generated in higher cortical areas, could serve this purpose. That is, top-down modulation might be the mechanism to induce perceptual learning (Caras and Sanes 2017). This perceptual learning might be implemented as modifications in the synaptic connections of the primary sensory circuitry. Another possible explanation that would not require reward-based learning, would be provided from a predictive coding perspective in which higher cortical areas provide a prediction signal observed during the late evoked response (Rao and Ballard 1999; Friston 2018). From this perspective, the repeated stimulation would shape such predictions via sensory adaptation processes (Whitmire and Stanley 2016; Weber and Fairhall 2019). Further experiments to test these ideas are necessary.

### Organization of the evoked responses in different sensory modalities

When we compared the cortical evoked responses to different sensory stimulation, we found that the temporal organization of the evoked responses, although similar, are not exactly the same across sensory modalities (Mohajerani et al. 2013). In particular, as shown previously (Mohajerani et al. 2013), the stimulation of the forepaw and hindpaw causes a rapid early evoked response in the corresponding areas of the somatosensory cortex that is comparable to the response in auditory cortex (around 20 ms after stimulus onset) but significantly faster than the response in the visual cortex (around 50 ms after stimulus onset) (Fig. 1 D-E). There is no difference in the duration of the early evoked response across the hindlimb, forelimb, auditory and visual stimulation. In contrast, we found a difference in the duration of the late component of the evoked response for different sensory modalities (Fig. 1E). These results indicate that although the two-component temporal organization of sensory evoked responses exists across modalities, the early evoked response is similar in duration but different in its response time. In contrast, the late component is similar in response time but different in duration.

### Experience dependent changes in the sensory evoked response and their functional role

In our experiments we observed that after repeated stimulation, there is an increase in the amplitude of the late component of the sensory evoked response which enhances organization of the evoked response into two components (Fig. S1 A), quantified as an increase in the slope of the late evoked component (Fig. S1 C), as an increase in the fit of a two gaussian model (Fig. S1 D-E). This modification lasts up to two hours after repeated stimulation (Fig. 5). Our study extends similar findings from rat visual system (Han et al. 2008), by examining the patterns over much larger regions of brain. This reverberation of the stimulation-induced pattern during subsequent spontaneous activity resembles the same timescale of reverberation reported in temporal patterns of cortical and hippocampal single-unit activity in rats (Euston et al. 2007; Bermudez Contreras, Gomez Palacio Schjetnan, et al. 2013). These results provide evidence that these sensory induced changes share some of the characteristics of synaptic plasticity mechanisms observed during memory formation processes (Bermudez Contreras, Gomez Palacio Schjetnan, et al. 2013). These results suggest that the repeated stimulation increases synaptic strengths in the brain circuitry involved in the early and late evoked responses. However, additional experiments are needed (e.g. NMDA blocking) to test this hypothesis.

Why would the late evoked response increase after repeated stimulation? According to recent research, animals that are better learners in sensory discrimination tasks (visual or tactile) also show an increase in the late evoked response compared to animals that do not learn the task (Sachidhanandam et al. 2013; Funayama et al. 2015a; Manita et al. 2015a). In our study, we observe this increase in the membrane depolarization (measured with voltage imaging) and synaptic activity (measured with glutamate imaging) signal at the mesoscopic level on average (Figs. 2, 5). This increase could be due to different mechanisms such as sensory adaptation (Solomon and Kohn 2014; Whitmire and Stanley 2016). For example, it could be due to physiological or morphological neuronal changes (e.g. lowering spiking threshold) or changes at population level such as an increase in the recruited neuronal population due to increases in the connectivity between the neurons that generate the late evoked response and the primary sensory neuronal population from which we are recording. All these are complex processes and require more experiments to be investigated.

This leads to the next question of why the late evoked response not only increases in amplitude, but also reorganizes such that the spatiotemporal pattern of the late evoked response becomes more similar to the one of the early evoked response. Since we did not address this directly, we can only provide a speculative answer. The reorganization of the sensory-evoked responses observed after repeated stimulation, likely involving cortical plasticity (Bermudez Contreras, Gomez Palacio Schjetnan, et al. 2013), causes a circuitry refinement in which a basin of attraction is formed by repeated stimulation. This idea has been used to explain the variability in the activity in sensory cortex (Hennequin et al. 2018). Therefore, subsequent incoming activity to the primary sensory areas, such as the late evoked response, falls into this basin of attraction and the pattern of the early evoked response is replayed. Whether this is an epiphenomenon of the cortical circuitry and plasticity mechanisms remains unknown and requires further experiments in which sensory perception is necessary to perform a behavioural task.

### Limitations and Future work

Different avenues are left unexplored in this work such as the relationship between cortical state and the organization of the evoked response. It is known that the trial-to-trial variability of sensory evoked responses is dependent on cortical state (Marguet and Harris 2011; Pachitariu et al. 2015; Schölvinck et al. 2015). Similarly, the early-late component organization of the sensory evoked responses have been proposed to depend on cortical state. According to (Curto et al. 2009), a bimodal structure of the evoked response can be largely explained by the cortical state preceding the stimulus presentation. Although in our recordings we confirmed that the brain state was more desynchronized after amphetamine injection, and that the activated the brain state remained stable for at least one hour (Fig. S3), our trial-to-trial analysis of the organization of the evoked response did not show a relationship with the cortical state preceding the sensory stimulation. At the behavioral level, it has been shown that the cortical state preceding the stimulus presentation did not affect the performance in a simple sensory perception task in head-restrained mice (Sachidhanandam et al. 2013). A deeper understanding at the single trial level in a preparation in which the cortical state could be quantified and separated from behavior, such as the one presented here, could shed some light on this issue. On a similar note, with the approach presented here, it should be possible to study the relationship between the changes observed during the late evoked response and other higher cortical regions (Crochet and Petersen 2015; Manita et al. 2015b). For example, an analysis of the activity propagation could reveal whether a cortico-cortical connectivity could explain the increase in similarity between the early and late components at the single-trial level.

Finally, even though we show that repeated stimulation given to awake animals increases the similarity between the spatiotemporal patterns of activity during the early and late components of the evoked response in auditory cortex (Fig. 6), we believe that the perceptual content of this signal might be limited. We know that perception might be affected by cognitive processes such as attention and motivation (Corbetta and Shulman 2002). Since the animals in our experiments were passively listening to tones, we believe that changes in the late evoked responses might be enhanced when the sensory stimulation has ‘meaning’ for the animal. Future experiments to test this would need to include a sensory-discrimination task in which sensory perception is directly associated to task performance.

### Significance

The results presented in this work demonstrate how the late evoked response that is usually associated with perception in sensory cortices is modified by experience at the mesoscale level. This modification consists in a spatiotemporal reorganization of the early and late components of the evoked response becoming more similar to each other. The reported spatiotemporal changes of the late sensory evoked response induced by repeated stimulation occur in different sensory modalities and last up to one hour. This work expands on previous studies of the structure of evoked cortical responses to sensory stimulation and opens important questions on the dynamics, perceptual content of the late evoked response and its relationship with experience. Future awake experiments involving perceptual tasks will help to expand on the understanding of the brain mechanisms involved in these phenomena.

## Acknowledgments

The present review was supported by Canadian Institutes of Health Research (CIHR) Grant# 390930 (MHM) and #199179 (AL), Natural Sciences and Engineering Research Council of Canada (NSERC) Discovery Grant #40352 (MHM) and #04636 (AL), Alberta Innovates (CAIP Chair) Grant #43568, Alberta Alzheimer Research Program Grant # PAZ15010 and PAZ17010, and Alzheimer Society of Canada Grant# 43674 to MHM. We thank Jianjun Sun for assistance with surgeries, Behroo Mirza Agha and Di Shao for animal husbandry. We thank Surjeet Singh for help with the image registration procedure. We thank Rasa Gulbinaite for feedback on this manuscript.

## Author contribution

E.B.C., A.L. and M.H.M. designed the experiments. E.B.C. and A.G.P. conducted the experiments. E.B.C. performed data analysis. E.B.C and M.H.M. wrote the manuscript, which all authors commented on and edited. M.H.M. supervised the project.

## Supplementary material

**Fig. S1.**
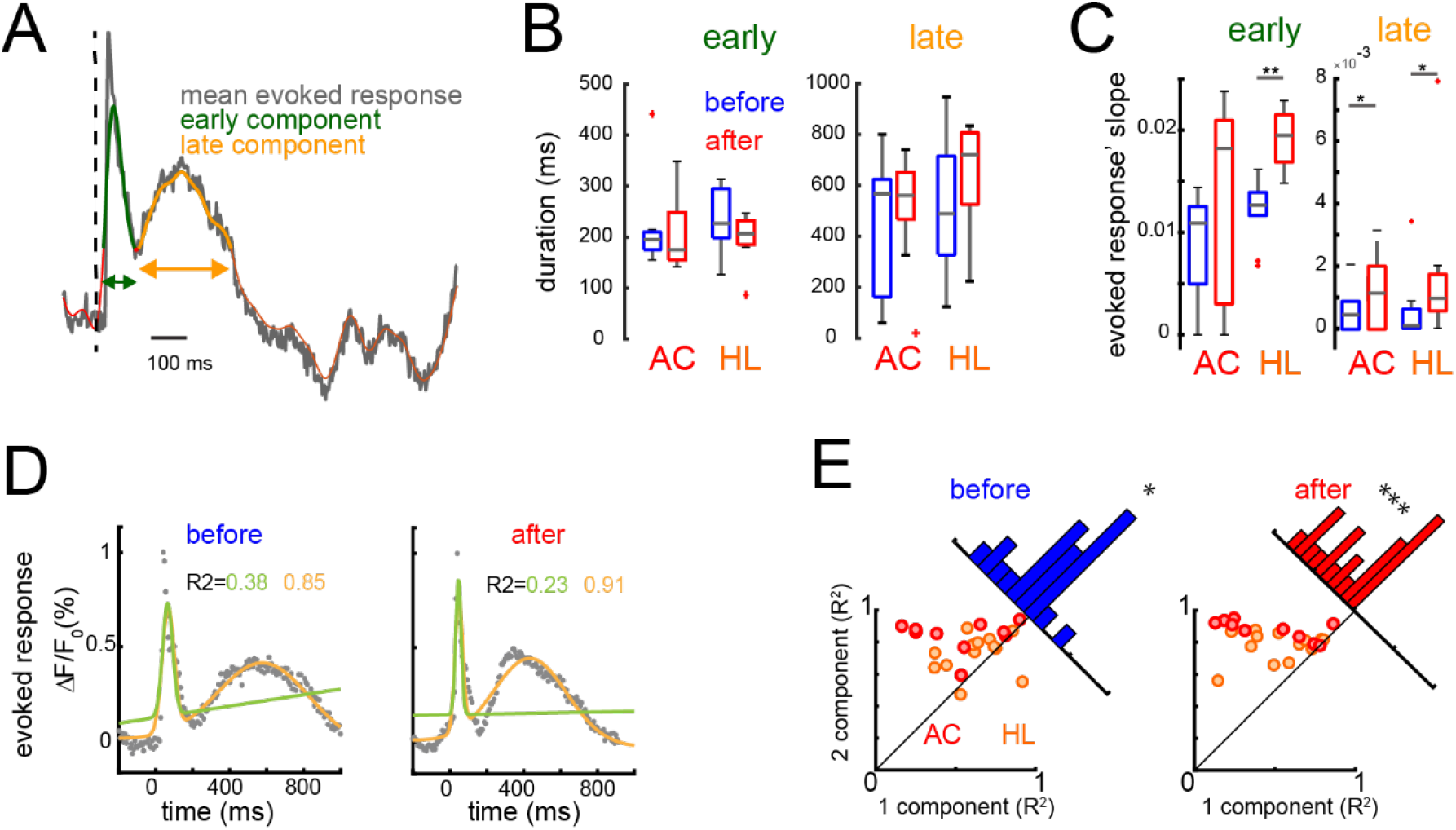
Organization of the evoked responses. **(A)** Sensory evoked responses are organized into an early (green) and late (yellow) components. **(B)** Box plots of the duration of the early (left pane) and late components (right pane) of auditory evoked stimulation (AC) and hindpaw stimulation (HL) before (blue) and after (red) repeated auditory and hindpaw sensory stimulation respectively. **(C)** Similar to B but for the slope of the early (left) and late (right) evoked components for the evoked responses to auditory stimulation (AC) and hindpaw stimulation (HL) before (blue) and after (red) repeated stimulation. Stars denote significant differences between the populations (p<0.05, p<0.01, Wilcoxon rank sum test or t-test depending whether the distributions were normal or not). **(D)** Example of approximation of single-trial evoked responses using a Gaussian Mixture Model with one (green) or two (yellow) components before (left) and after (red) repeated stimulation **(E)** Scatter plots of R-squared using one or two components before (blue) and after (red) repeated auditory or hindpaw stimulation (respectively). Each dot denotes the mean R-squared for the corresponding model (1 or 2 components) across trials of each animal for hindpaw (orange) or auditory (red) stimulation. The stars represent p<0.05 and p<0.001, t-test).

**Figure S2.**
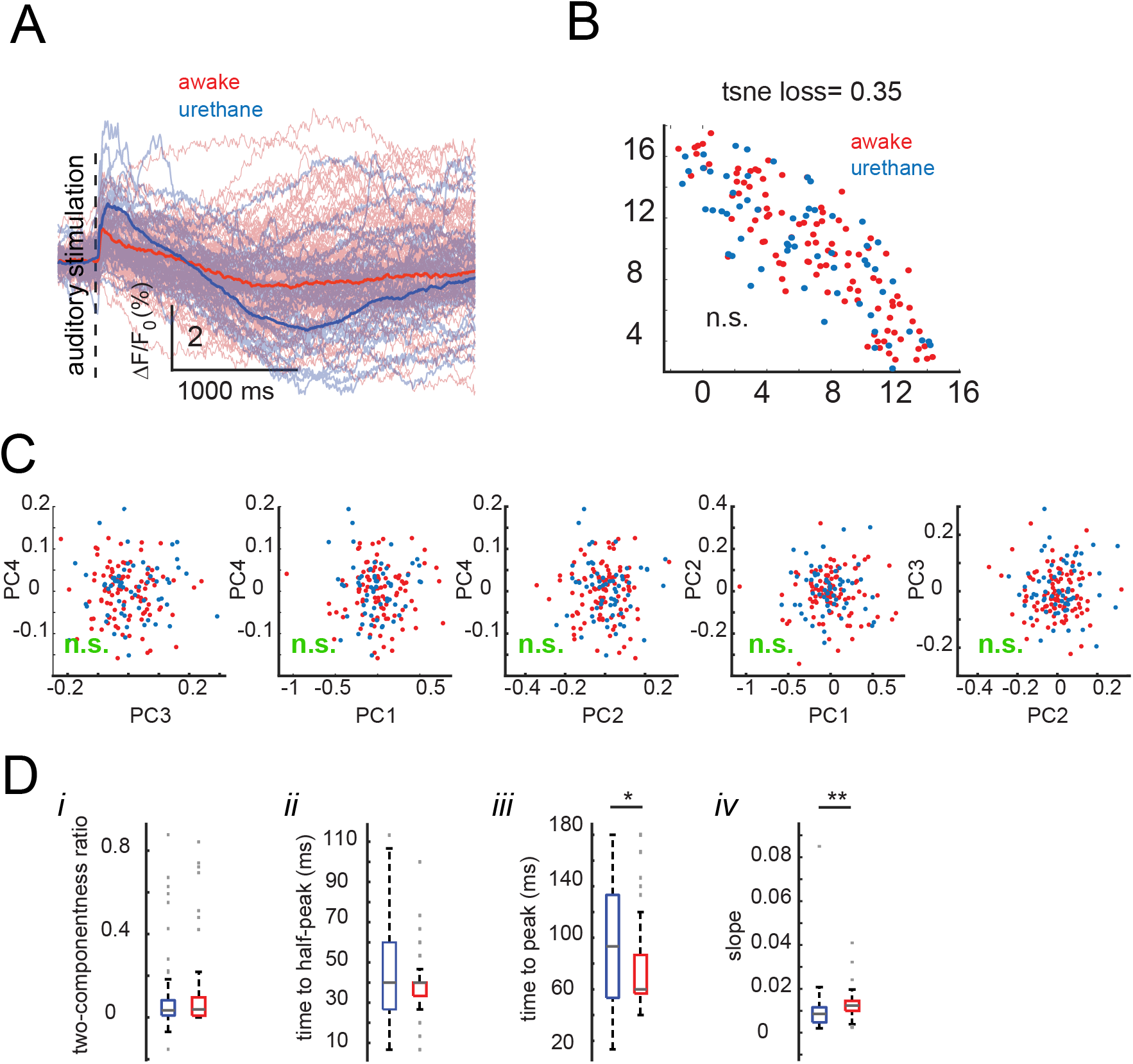
Sensory evoked activity in awake and anesthetized animals. **(A)** Tone-evoked responses in the auditory cortex in awake (red) and anesthetized (blue) mice (n=60 trials) thick traces correspond to the mean across trials, thin traces correspond to the evoked trials. **(B)** T-SNE projection of single-trial evoked responses during wakefulness (red) and urethane-anesthesia (blue). **(C)** Comparison of the four principal components of single-trial evoked responses (n. s. denotes not significantly different correlation coefficient using the Fisher’s z transformation). **(D)** Comparison of the early evoked responses in awake (red) and urethane anesthetized (blue) mice. (i) Two componentness (mixture of Gaussian model). (ii) time-to-half-peak (iii) time-to-peak. (iv) Slope of single-trial evoked responses from stimulus onset to peak.

**Figure S3.**
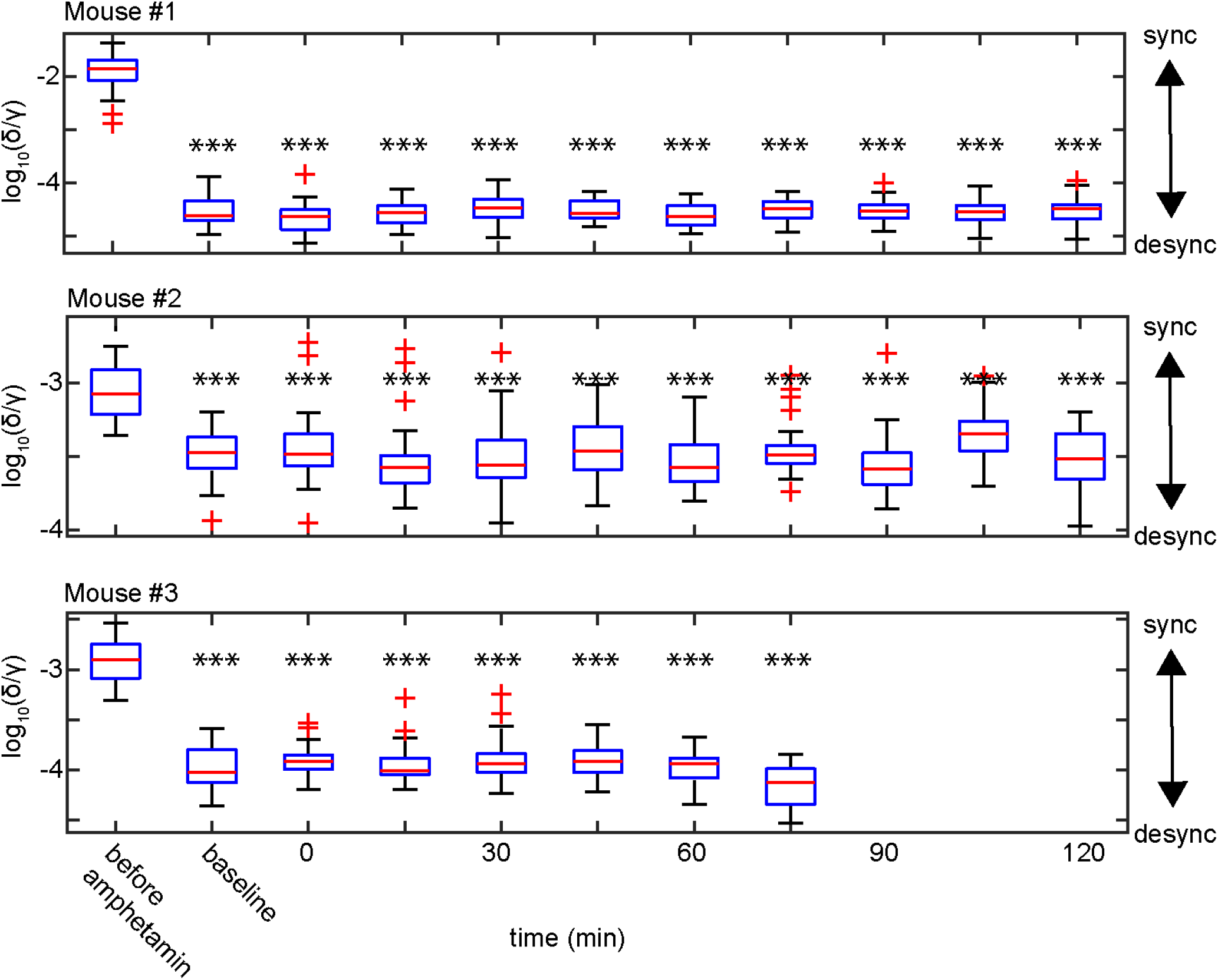
Cortical state stability. Each panel shows the brain state for separate animals across the corresponding recordings. The brain state before amphetamine was injected was significantly different from every recording for up to two hours (one-way ANOVA, Bonferroni correction, *** represents p<0.001). In contrast, the brain state was not significantly different across recordings after amphetamine injection.

**Figure S4.**
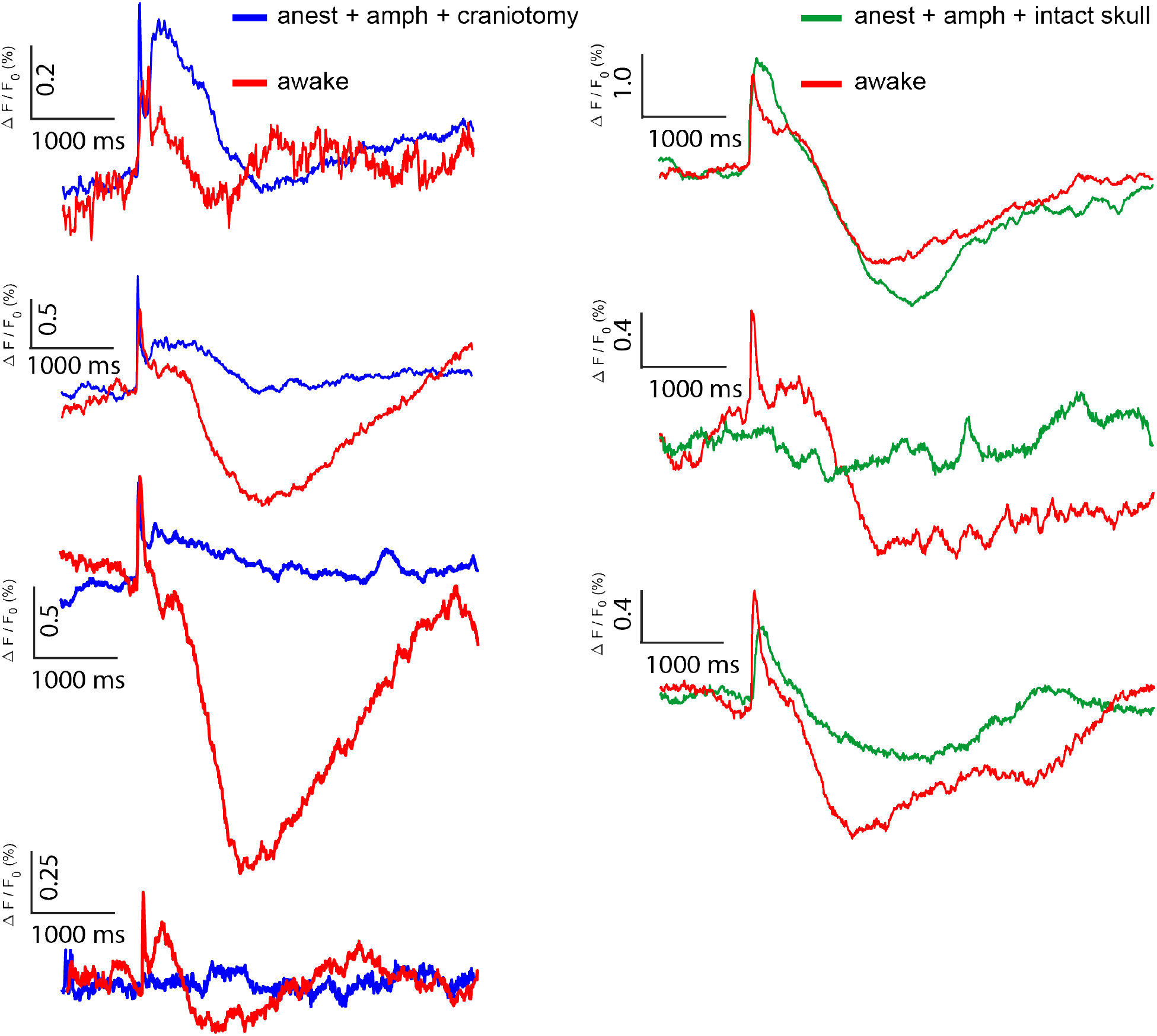
Average evoked responses in awake and anesthetized animals. The evoked responses to a single tone stimulation were recorded in awake or anesthetized (blue) animals with a cranial window (red, left column) and also in anesthetized (green) with skull intact or awake (red, right column) preparations. Each trace represents the average evoked response over trials (n=30 trials) for seven animals.

